# A Neurostimulator System for Real, Sham, and Multi-Target Transcranial Magnetic Stimulation

**DOI:** 10.1101/2021.11.22.469567

**Authors:** Majid Memarian Sorkhabi, Timothy Denison

## Abstract

**Background:** Transcranial magnetic stimulation (TMS) is a clinically effective therapeutic instrument used to modulate neural activity. Despite three decades of research, two challenging issues remain, the possibility of changing the 1) stimulated spot and 2) stimulation type (real or sham) without physically moving the coil.

**Objective:** In this study, a second-generation programmable TMS (pTMS2) device with advanced stimulus shaping is introduced that uses a 5-level cascaded H-bridge inverter and phase-shifted pulse-width modulation (PWM). The principal idea of this research is to obtain real, sham, and multi-locus stimulation with the same TMS system.

**Methods:** We propose a two-channel modulation-based magnetic pulse generator and a novel coil arrangement, consisting of two circular coils with a physical distance of 20 mm between the coils and a control method for modifying the effective stimulus intensity, which leads to the live steerability of the location and type of stimulation.

**Results:** Based on the measured system performance, the stimulation profile can be steered ± 20 mm along a line from the centroid of the coil locations by modifying the modulation index.

**Conclusion:** The proposed system supports electronic control of the stimulation spot without physical coil movement, resulting in tunable modulation of targets, which is a crucial step towards automated TMS machines.

## 1. Introduction

Transcranial magnetic stimulation (TMS) is a non-invasive technique used to stimulate and modulate cortical neurons [1] [2]. TMS is based on the fundamental principles of electromagnetic induction: a brief, strong electric current is delivered to the windings of a coil, inducing a changing magnetic field, which in turn induces an electric field in the cortex. Applying magnetic stimulation to neural cells can depolarize and hyperpolarize neurons. TMS is commonly used as a diagnostic tool for various neurological disorders; it is an FDA-approved treatment of, for example, major depressive disorder, and is being considered as a potential therapy for many other applications [2] [3] [4].

Conventional magnetic stimulators consist of three main elements: a power capacitor, an inductor acting as the stimulation coil, and a thyristor switch to control the connection between them [5]. Most stimulators are restricted to specific stimulus frequencies and shapes, generating only monophasic or biphasic cosine-shaped pulses. Typically, the stimulus frequency is approximately 2.5 kHz, but the exact value is determined by the architecture of the stimulator and the coil. During biphasic stimulation, some of the energy delivered by the pulse is recovered, enabling the generation of up to 100 pulses per second [6]. However, in monophasic stimulation, the energy is dissipated through a resistor [5], only allowing stimulation patterns of one pulse per second at full power.

The ancillary effects of TMS, such as clicking noise, stimulation of nearby peripheral nerves, and scalp and facial sensations may interfere with task performance in clinical research through subject biasing and distraction, which can contaminate trial outcomes [7] [8]. Subjects in the control group, who were exposed to all artifacts but did not receive stimulation in the target brain area, could help to separate the effects of the auditory and tactile artifacts from the test group. Physically tilting the coil from 45° to 90° from the target brain area, electrical scalp stimulation during the sham TMS [9], flipping the current direction in each of the Figure-of-eight coil coils by an external custom-made switch box or changing the direction of the winding inside the coil, are examples of conventional methods for implementing a sham-TMS protocol [10] [11].

Another constraint is that maintaining the target location of stimulation is a considerable challenge of the TMS technique. Neuro-navigation technology can sustain the target in a couple of millimeters using robotic arms and physical coil displacement [12]. In addition to the high cost and complexity of these systems, the safe handling of heavy coils is relatively slow, and it is practically impossible to move the target location during rapid stimulation protocols. Using an array of coils, such as a 16-coil or 5-coil, can theoretically solve the problem of moving the target location, but a separate TMS system is required to drive each coil [13]. In addition to being bulky and expensive systems, the dynamics of conventional pulse generators will probably not allow the stimulus parameters to be modified quickly enough for repetitive protocols. The multi-locus TMS system introduced by Koponen et al. [13], which consists of an optimized figure-of-eight coil and an oval coil stacked on top of each other, can move the stimulated spot in a ±15 mm long line from the coil center, which has certain limitations in terms of the speed of spot movement and stimulus generation for rapid TMS protocols.

Recently, the use of isolated-gate bipolar transistors (IGBTs) instead of thyristors, as well as the implementation of H-bridge structures, has enabled more control over the stimulation parameters. Peterchev et al. developed a series of transcranial magnetic stimulators with controllable pulse parameters (cTMS) [14] [15] [16] that produce both monophasic and biphasic stimuli of different frequencies and allow more pulses per second than conventional TMS devices. However, the high current stress on the IGBTs and the limited number of pulses that can be selected limit the achievable protocols [17].

A new approach using pulse width modulation (PWM) enables the imitation of any arbitrary stimulus while reducing the current stress on the IGBTs, called programmable TMS or pTMS [17]. The proposed structure can reduce the current stress on the power switches by paralleling IGBTs by recovering the energy delivered to the coil, which can generate magnetic pulses with a high repetition rate (up to 1 kHz), low interstimulus intervals (1 ms), and low voltage decay. Preliminary results of in-human experiments reveal that the conventional monophasic and its PWM equivalent pulses can induce almost identical results on the primary motor cortex, and the PWM stimuli appear to fundamentally mimic the effect of conventional magnetic pulses [18].

Nowadays, power converter technology has been applied in many fields, such as motor drivers, electric vehicles, renewable energy sources, and magnetic resonance imaging (MRI) scanners. Among power converter methods, multilevel inverter topologies, including cascaded H-bridges (CHB), flying capacitors, and modular multilevel converters (MMCs), are industrially accepted solutions for medium-voltage high-power applications. Broadly, the CHB architecture has received more attention because of its simple layout and construction, high modularity, uncomplicated control of power switches, the need for fewer power switches, and no voltage balance problems in energy storage capacitors. These features make cascaded H-bridge inverters a convenient solution to be applied as a TMS pulse generator core. This study introduces the second generation of a programmable TMS device that uses these advatages with the goal of generating arbitrary stimulus shapes via the PWM technique.

This research also aims to develop a versatile TMS system, including a pulse generator and a novel coil arrangement, to control the stimulation type and location. To measure the performance of the proposed system, we built a two-channel pTMS device and a non-overlapping two circular coil.

## 2. Materials and Methods

### 2.1 Coil design

A custom-made figure-of-eight coil with two loops of 70-mm-diameter circular coils, where the distance from the edge to the edge of the two coils is 20 mm, and each of the two wings is controlled separately was fabricated (Magstim Company Ltd, UK). Both coils were placed individually in a coil enclosure. The coils were air-cooled and had four independent temperature monitoring points on the coils. The left and right coil inductances were measured at 18 μH and 18.3μH, respectively, and because of the physical distance between the coils, their mutual inductance can be neglected. More details are available in the appendix (Figure A.1).

Electromagnetic simulations were performed using the COMSOL software (V5.3, Sweden) with the finite element method (FEM), using the AC/DC module and the stationary solver. A 3D MRI-derived head model was used to calculate the induced electric field (E-field). The head tissue was assumed to be homogeneous, with an electrical conductivity of σ= 0.333 S/m [19] [20]. For FEM analysis, the final mesh structure of the coil and head comprised 145,000 nodes and 3.1 million tetrahedral elements, with a minimum element size of 5 μm. A linear solver with a relative error of 1.2e-6 was selected. During the FEM simulation, the coil and head models were in the air [21].

To validate the modeling results and estimate the accuracy of the pulse generator and the manufactured coil, a custom system with a pickup coil was designed that swept the 40 × 40 mm area with 5 mm steps and measured the induced E-field.

### 2.2 Magnetic pulse generator

As shown in Figure 1, the CHB inverter consists of a series of N power cells, usually based on identical H-bridge sub-modules that are cascaded from the cell output. Each of them includes a galvanically isolated DC link as a voltage source and an energy storage capacitor. If the DC link voltage is equal in all the cells, CHB is classified as a symmetrical inverter, but if the voltages of each power cell are dissimilar, it is classified as an asymmetrical architecture. Although an asymmetrical CHB with a smaller number of power cells can generate a greater voltage level, controlling the returning (regenerative) currents from inductive loads, such as stimulation coils in TMS devices, and maintaining capacitor voltage balance can be challenging. In addition, modularity and lower implementation and maintenance costs are other advantages of a symmetrical CHB over an asymmetrical inverter [22]. In this study, a symmetrical structure was selected and implemented. Each H-bridge can be used to create three output voltage levels: {-V_DC_, 0, +V_DC_}. Thus, by connecting two cells, five voltage levels {-2V_DC_, -V_DC_, 0, +V_DC_, +2V_DC_} can be achieved for the output pulse. As represented in Figure 1(c), the output voltages of the H-bridges (V_1_(t) and V_2_(t)) are similar but displaced in time, and the total output pulse (V_out_(t)) is the sum of the outputs of the two cells.

**Figure 1.**
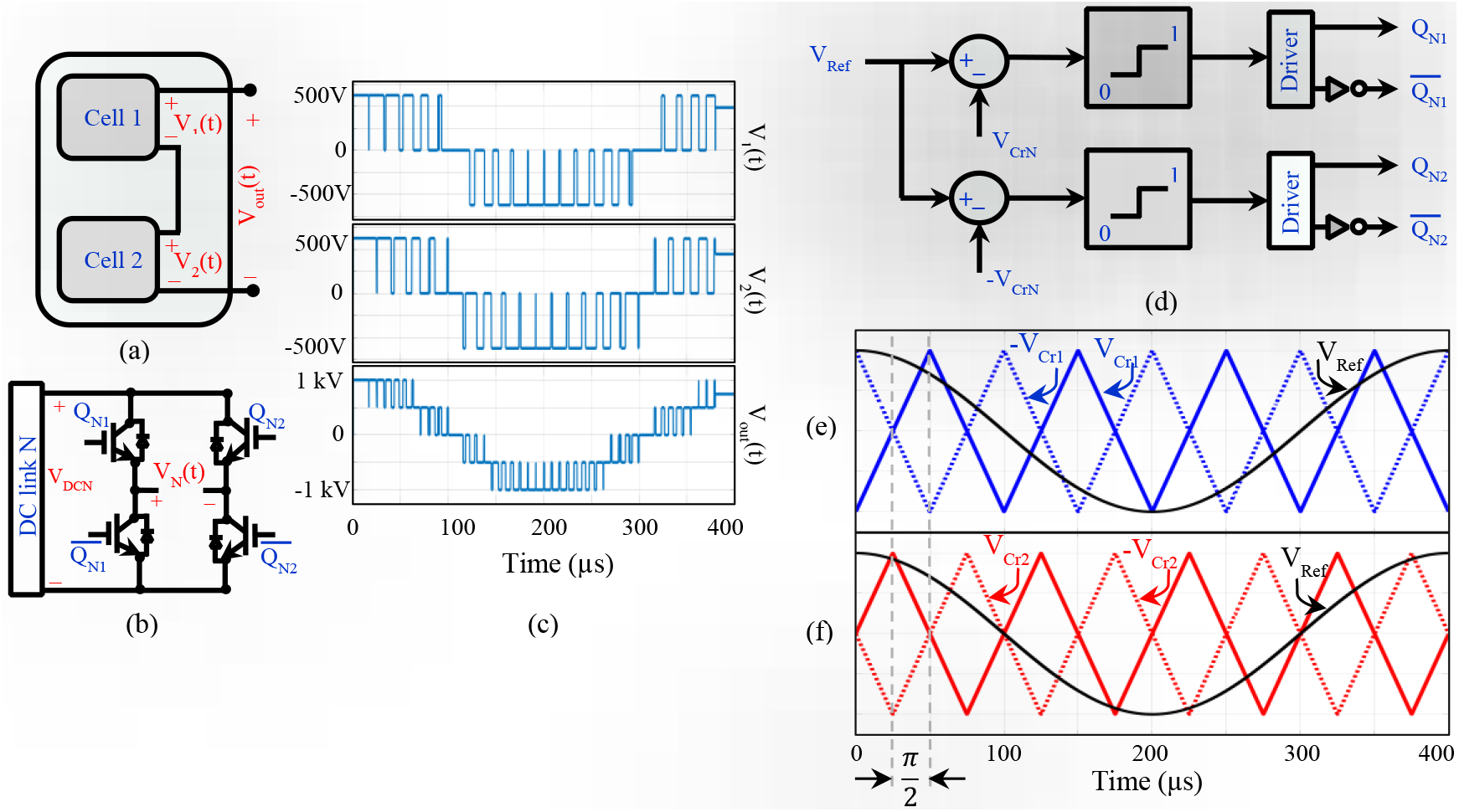
Single-phase two-cell cascaded H-bridge power converter and control method. (a): Topology of power cell connection. (b): H-bridge structure, where N represents the cell number. (c): Output voltages of power cells and total output voltage using pulse width modulation. As an example, the voltage level of each DC link is 500 V and the output pulse is a 5-level 2.5 kHz cosine-shaped equivalent modulated signal. (d) Logic block diagram to generate the control signal, where V_Ref_, V_Cr_, and N represent the reference stimulus, triangular carrier pulse and cell number, respectively. (e)-(f) The phase-shifting technique uses a cosine wave as a reference signal (e) for controlling the first cell switches and (f) for controlling the second cell switches.

For a multilevel inverter, multicarrier phase-shifted PWM (PS-PWM) is a common method for creating the switching pulses of the power devices [23] [24] introduced in Figure 1 (d). In PS-PWM, triangular carrier pulses (V_cr_) with the same frequency but shifted by a specific phase are compared with the desired reference signal (V_Ref_) to determine the trigger signal for the power switches 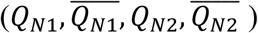. The carriers are shifted by π/N rad, where N is the total number of H-bridges. Figure 1(d-f) shows the PS-PWM method for the structure implemented in this study, a two-cell CHB inverter where carriers have a phase difference of π/2 rad [25] [26].

The PS-PWM method has a multiplicative effect on the switching frequency. While each cell can be triggered at a low switching frequency, the final output pulse is the sum of the output voltages of the cells (V_1_(t)+ V_2_(t)= V_out_(t)) and its harmonic components will be at higher frequencies than the output of each cell [27]. As an example, if the PS-PWM technique is employed in an N-cell CHB and the triangular carrier frequencies are F_PWM_, due to the switching, the first harmonic in the output voltage (V_out_(t)) will be located at N*2*F_PWM_ [25] [28]. This feature makes the CHB inverter very attractive for TMS pulse generators. Because the inherent low-frequency property of the neural tissue is expected to eliminate the high-frequency harmonics produced by switching, the neurons would only respond to the main harmonic component, which is the reference pulse frequency.

In this study, a two-channel TMS device was used to generate magnetic stimuli. For this purpose, for each coil channel, two full H-bridges are cascaded together, which can generate 5-level PWM pulses, as shown in Figure 2. The pulse capacitors of each H-bridge are implemented separately, but two common isolated DC sources (including step-up transformers and rectifiers) are used for both channels. Parallel-connected IGBTs are used to allow high coil currents to be achieved (three IGBT modules per leg) [29] [30]. The DC link voltage is 1.5 kV (750 V per H-bridge) and the measured peak current for the 2.5 kHz biphasic pulse is 6 kA.

**Figure 2.**
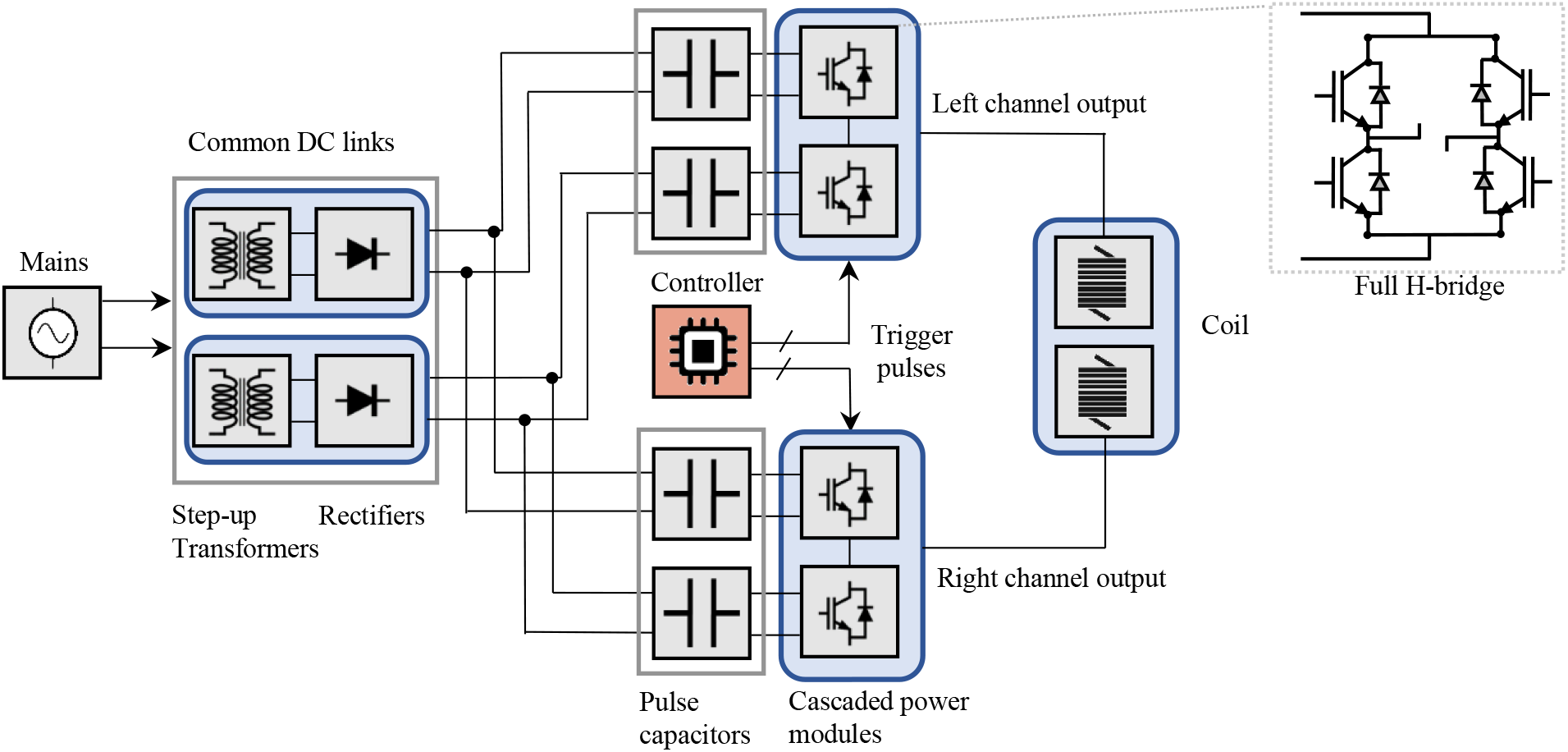
The overall diagram of the proposed two-channel TMS system, which is built by cascading two full H-bridges in each coil channel. Key components of the proposed pulse generator: DC links include two isolated step-up transformers (custom manufactured, secondary voltage (RMS): 570 Vac, 5.7 kW, class-E insulation, Eastern Transformers, UK), two full-bridge diode rectifiers (STTH9012TV1, STMicroelectronics). Pulse capacitors consist of a series of two capacitors (10000 μF, 500 V_DC_, ALS70A103NT500, KEMET Electronics, EU), a total of 8 capacitors. The full H-bridges are built with three IGBTs in parallel (V_CEmax_= 1.2 kV, I_CRM_ = 1.8 kA, SEMiX603GB12E4p Semikron, Germany), for a total of 6 modules per H-bridge and 2 H-bridges per channel. Generation of trigger pulses for IGBTs and modulation index changes are done by a digital controller (Micro Lab Box, Dual Core 2 GHz processor, DS1202, DS1302 I/O, dSPACE, Germany).

The proportional-integral (PI) control loop mechanism is utilized in the CHB inverter control feedback, where the reference coil current is defined as the desired setpoint for the minimization of current distortions [31]. Parasitic resistance loss in the coil and switching losses of IGBTs are the main causes of these distortions in the TMS system, which can be minimized by optimizing the PWM patterns. This optimization method was calculated offline and a look-up table that holds the trigger pulses was generated. The content of this look-up table is a function of the desired stimulus waveforms and treatment procedures. The offline computation was performed by modeling and reconstructing the circuit behavior utilizing the commutated current, voltage, and semiconductor characteristics. Finally, the controller parameters were tuned using a laboratory test bed. Moreover, the safe operating limits of the TMS machine were added to the controller design formulation. Even more imperative, for the PI controller, a significant part of the control effort is shifted from the hardware stage to the computational stage.

The filtering behavior of the neuronal membrane, when exposed to electromagnetic waves, causes the net effect of the applied pulse on the neuron to be a filtered pulse with a cut-off frequency of approximately 1 kHz [14]. Therefore, the applied staircase PWM pulses, which have high-frequency harmonics, are observed as a softened pulse in the neuron. The constant DC voltage in pulse capacitors and applied stimulation intensity alternation using the concept of amplitude modulation index greatly reduce the design complexity of AC to DC converters.

### 2.3 Amplitude modulation index

The concept of the amplitude modulation index (m) in multilevel inverters controls the effective intensity of the output pulse and provides an instant flexibility of the stimulation type and location by changing the pulse waveform. The modulation index, also known as the amplitude modulation ratio, is the ratio of the target pulse amplitude to the carrier signal amplitude, which is adjustable in the controller [32]. The CHB can produce an average output voltage that is linearly proportional to the target wave in the range of −1 ≤ m ≤ 1 [17]. Potential distortions are minimized with the PI controller as a result of using this concept to control the effective stimulation intensity [33].

## 3. RESULTS

Figure 3 shows the computational results for the induced E-field 20 mm below the coils, as the hypothetical location of the cortex, for the Magstim 70-mm Figure-of eight coil (P/N 3190)(Figure 3a) and the proposed coil)(Figure 3b). The E-field profile for real stimulations is similar for the proposed and Figure-of-eight coils, and the difference is in the lower field concentration at the overlap area.

**Figure 3.**
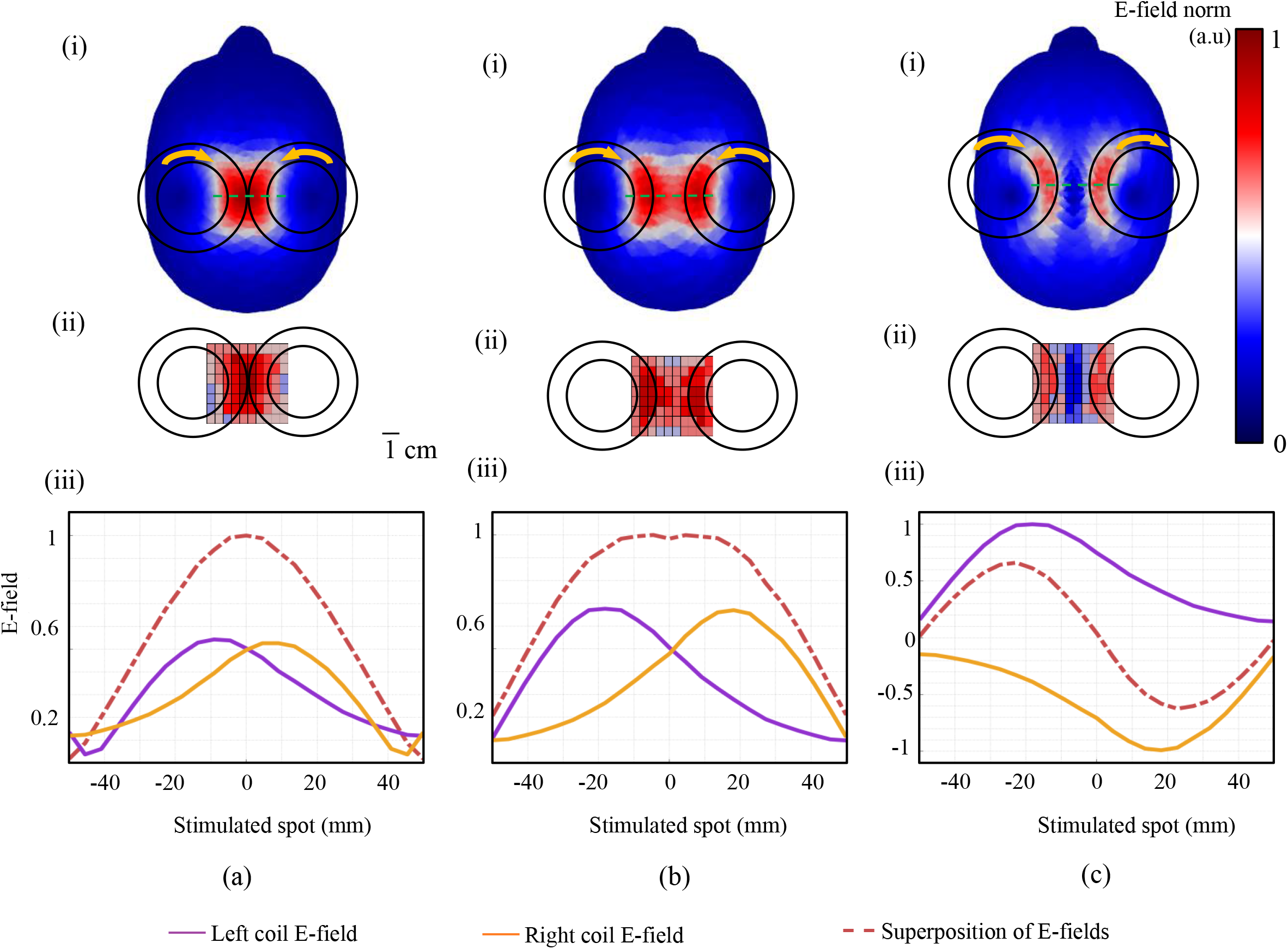
Concept of two independent circular coils, and validation of real/sham stimulations through both computational modeling and a pickup coil measurement. (a) Results for a Magstim 70 mm figure-8 coil, showing (i) color map of the spatial distribution of the induced E-field norm (ii) Induced E-field to the pickup coil with 5 mm steps in the target area and (iii) the E-field measured on the dashed green path shown in (i), for injecting or not injecting the stimulation current into the coils independently. (b) Results for the proposed system when the current direction in the coils is set according to the real stimulation (m_left_= m_right_= 1). (c) Results for the prototype when the current direction in the coils is set according to the sham stimulation (m_left_=1, m_right_= −1). In (iii), the solid purple curves 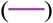 show the induced E-field when the stimulation current is injected only into the left coil and the solid orange curves 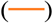 represents the E-field only by injecting current into the right coil and the dashed curves represent the induced field with the presence of current in both coils. All E-field measurements in (iii) were taken along the green dashed line shown (i) and E-field amplitudes were normalized to the maximum. In (i), the arrows on the coils indicate the direction of the electric current. The E-field norm means the amplitude of the electric field 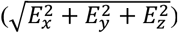.

For the sham stimulation (Figure 3c), it is obvious that in the central region, the E-fields of the two coils attenuate each other and no stimulation will occur, but in areas below the coils, it is possible to see the stimulation. This off-target stimulation can lead to a sensation of stimulation in the subject and potentially make the placebo effect more realistic. Measurements of sound pressure level for the same pulse intensities were not statistically different between the real and sham stimulation (~70 dB at 80% of the maximum device output, p = 0.45).

The results of the pickup coil measurements are shown in Figure 3(ii); by comparing them with the computational modeling outcomes, it can be concluded that the performance of the implemented system is similar to that of the modeling. To investigate the effect of each coil loop on the final E-field profile, the coils were driven separately, and the peak E-fields were measured, as shown in Figure 3 (iii). Based on the results, the final stimulating field is the sum of the fields from each coil; therefore, by appropriately modifying the effective pulse intensity (modulation index), the induced E-field can be shaped and targeted without physical movement of the coils themselves.

The physical translation of the stimulated spot while holding the locations of the coils constant was explored and illustrated in Figure 4 (a). According to the measurement results for 21 different modulation indexes, by changing the index, the stimulation profile can be steered ± 20 mm from the centroid of the coil locations in 2 mm steps. Three examples of monophasic PWM-stimuli generated by the implemented magnetic pulse generator are shown in Figure 4 (b)–(d), as well as the measured E-field at the center and ± 20 mm away from the center, to test the efficiency of the proposed system (i-iii).

**Figure 4.**
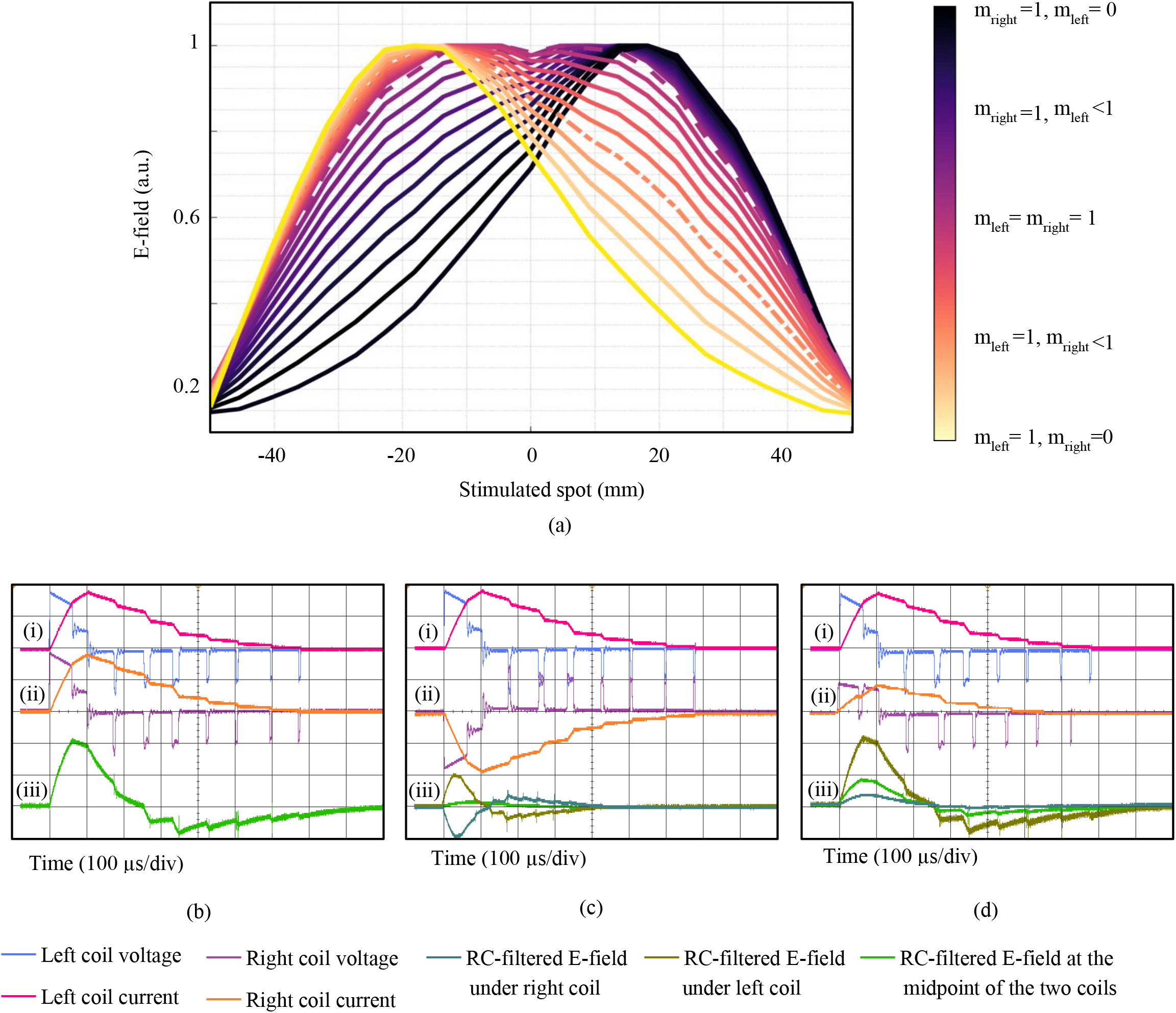
Ability to steer the stimulated spot by varying the modulation index. (a) The experimentally measured E-field distribution of the different modulation indexes along the green dashed line shown in Figure 3(i). For each measurement, one index was kept constant at 1 and the other index was changed in steps of 0.1 (0-1), for a total of 21 different indexes. Measured voltage and current waveforms for (b) m_left_= m_right_= 1, (c) m_left_=1, m_right_= −1, (d) m_left_=1, m_right_= 0.4, where (i) presents left-coil voltage 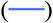 and current 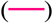, (ii) presents right-coil voltage 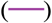 and current 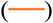 and (iii) represents the measured E-field after passing through an RC-lowpass filter, at the midpoint of the two coils 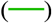, 2 cm to the left 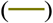 and 2 cm to the right of the center 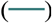, as an estimate of the neural membrane voltage change after stimulation. The TMS-induced E-fields generated by the pulses of (b) and (d) are shown as dashed and dash-dotted lines in (a), respectively. The E-field generated by the pulses of (c) is also presented in Figure 3(C-iii). In (i) and (ii), for the voltage and current graphs, all amplitudes were normalized to the maximum. In all time axes, each square represents 100 μs. All measurements were performed with a digital oscilloscope at a sampling rate of 500 MSa/s. No bandwidth limitations or filters were employed to remove the switching glitches/spikes.

The coil voltages and currents were measured using a high-voltage differential probe (TA044, Pico Technology, UK) and a Rogowski current probe (I6000S FLEX-24, Fluke, USA), respectively. The peak coil current and delivered energy were measured to be 6 kA and 250 J, respectively. The electric field was measured using a pickup coil. The behavior of the nerve tissue in the presence of an external electric field is modeled with a passive RC filter with a time constant of 150 μs, as an accepted model for magnetic neuron stimulation [34] [17]. The conclusion is that changing the modulation index shifts the activation site.

## 4. Discussion

Power converters have been gaining popularity in TMS equipment owing to their potential to emulate arbitrary magnetic stimuli [16] [17]. The main reasons for the progress of the CHB are the reduction in power ratings and dv/dt across switches and the alleviation of the cost by reducing the power switch count. Owing to the modularity of CHB, it can be stacked for high-power magnetic stimulation applications. To generate a flexible magnetic pulse, cascaded H-bridge inverters must synthesize a staircase waveform using the PWM method. The phase-shifted PWM method applied to CHB inverters enables highly effective overall switching frequencies for the generated stimulus, even with a low switching frequency of each individual switch [26].

Nurmi et al. demonstrated that there exists a trade-off between the induced E-field focality and the number of coils stacked on top of each other [35]. Optimal coil arrangements can provide wide control over the cortical region, whereas each coil requires a separate pulse generator and increases the system complexity. The use of two optimized coils on top of each other, and high-precision capacitive chargers proposed by Koponen et al. [13], enables more focused targeting and shifting of the stimulated spot compared to this study. Because of the pulse generator circuit structure and elements used in it, such as capacitor chargers and relay-boards utilized for high-voltage sharing, it is not possible to charge/discharge the pulse capacitors quickly [36], so the stimulation location can be moved only in one direction and by increasing the voltage of the capacitors. In addition, the ability to change the stimulation type from real to sham has not been investigated in that structure.

As a limitation of the proposed system, owing to the physical distance of the coils, larger currents must be injected into the single coils to superimpose the fields of the two coils and achieve the stimulation profile of the Figure-of-eight coil, as shown in Figure 3 (iii).

## 5. Conclusion

The present neurostimulator establishes the unique potential of cascaded H-bridge inverter topologies to generate an arbitrary magnetic stimulus. The modular property of this inverter enables the improvement of the neuromodulation waveform by cascading H-bridges. The proposed pulse generator along with novel coil suggests the possibility of ‘‘live steering” of stimulation type and spot without having to physically move the coils themselves by modifying the effective intensity delivered to a fixed set of coils, as an example the coil can be driven in-phase to implement sham stimulation.

## Supporting information

Constricted coil for real, sham, and multi-target stimulation

## Acknowledgment

The authors would like to thank Magstim Company Ltd (UK) for construction, providing the stimulation coil, and valuable guidance on design considerations.

## Authorship contributions

MMS and TD conceptualized the study and drafted the manuscript. MMS performed the modeling, experimental measurements, and statistical analyses. TD provided the funding for this study. All authors have revised and approved the final version of the manuscript.

## Compliance with ethics guidelines

All authors declare that they have no conflicts of interest or financial conflicts to disclose.

## data availability statement

Data, and code used in this work are available in the public domain upon direct request from corresponding author.

## Appendix A. Supplementary data

Supplementary data to this article can be found.

## References

[1] B. J. Edelman and et al., “Systems Neuroengineering: Understanding and Interacting with the Brain,” Engineering, vol. 1, no. 3, pp. 292–308, 2015.

[2] J. P. Lefaucheur, N. André-Obadia, A. Antal, S. S. Ayache, C. Baeken, D. H. Benninger, R. M. Cantello, M. Cincotta and e. al, “Evidence-based guidelines on the therapeutic use of repetitive transcranial magnetic stimulation (rTMS),” Clinical Neurophysiology, vol. 125, no. 11, pp. 2150–2206, 2014.

[3] Y. Zhang and et al., “Identification of psychiatric disorder subtypes from functional connectivity patterns in resting-state electroencephalography,” Nature Biomedical Engineering, 2020.

[4] m. Memarian sorkhabi and et al., “Numerical modeling of plasticity induced by quadripulse stimulation,” IEEE Access, vol. 9, pp. 26484–26490, 2021/2/8.

[5] C. M. Epstein, “Electromagnetism,” in The Oxford Handbook of Transcranial Stimulation, E. Wassermann, C. Epstein, U. Ziemann, V. Walsh, T. Paus and S. Lisanby, Eds., Oxford, Oxford Universtiy Press, 2008, pp. 3–5.

[6] MAGSTIM company, “MAGSTIM RAPID2 P/N 3576-23-09 OPERATING MANUAL,,” The MAGSTIM Company LTD, 2009.

[7] E. M. Robertson, H. Théoret and A. Pascual-Leone, “Studies in Cognition: The Problems Solved and Created by Transcranial Magnetic Stimulation,” Journal of Cognitive Neuroscience, vol. 15, no. 7, p. 948–960, 2003.

[8] E. Grasin, I. Loginov, A. Masliukova and N. Smirnov, “Realistic sham TMS,” Brain Stimulation, vol. 12, no. 2, p. 418, 2019.

[9] M. Mennemeier and et al., “Sham Transcranial Magnetic Stimulation Using Electrical,” Brain stimulation, vol. 2, no. 3, p. 168–173, 2009.

[10] F. Hoeft and et al., “Electronically Switchable Sham Transcranial Magnetic Stimulation (TMS) System,” PLoS ONE, vol. 3, no. 4, 2008.

[11] J. Ruohonen and et al., “Coil design for real and sham transcranial magnetic stimulation,” IEEE Transactions on Biomedical Engineering, vol. 47, no. 2, pp. 145–148, 2000.

[12] S. Kantelhardt and et al., “Robot-assisted image-guided transcranial magnetic stimulation for somatotopic mapping of the motor cortex: a clinical pilot study,” Acta neurochirurgica, vol. 152, no. 2, pp. 333–343, 2010.

[13] L. M. Koponen, J. O. Nieminen and R. J. Ilmoniemi, “Multi-locus transcranial magnetic stimulation—theory and implementation,” Brain Stimulation, vol. 11, no. 4, pp. 849–855, 2018.

[14] A. V. Peterchev, R. Jalinous and S. H. Lisanby, “A transcranial magnetic stimulator inducing near-rectangular pulses with controllable pulse width (cTMS).,” IEEE TRANSACTIONS ON BIOMEDICAL ENGINEERING, vol. 55, no. 1, pp. 257–266, JANUARY 2008.

[15] A. V. Peterchev, D. L Murphy and S. H Lisanby, “Repetitive Transcranial Magnetic Stimulator with Controllable Pulse Parameters,” J Neural Eng., vol. 8, no. 3, p. 036016, 2011.

[16] A. V. Peterchev, K. D’Ostilio, J. C. Rothwell and D. L. Murphy, “Controllable pulse parameter transcranial magnetic stimulator with enhanced circuit topology and pulse shaping.,” Journal of neural engineering, vol. 22, no. 5, 2014.

[17] M. Memarian Sorkhabi and et al., “Programmable Transcranial Magnetic Stimulation-A Modulation Approach for the Generation of Controllable Magnetic Stimuli,” IEEE Transactions on Biomedical Engineering, vol. 68, no. 6, pp. 1847–1858, 2020.

[18] M. Memarian Sorkhabi and et al., “A digital transcranial magnetic stimulator for generating arbitrary pulse-shapes and patterns,” Brain Stimulation, vol. 14, no. 6, pp. 1613–1614, 2021.

[19] M. Memarian Sorkhabi, K. Wendt, M. T. Wilson and T. Denison, “Estimation of the Motor Threshold for Near-Rectangular Stimuli Using the Hodgkin–Huxley Model,” Computational Intelligence and Neuroscience, pp. 1–10, 2021.

[20] M. Memarian Sorkhabi, J. Frounchi, P. Shahabi and H. Veladi, “Deep-brain transcranial stimulation: A novel approach for high 3-D resolution,” IEEE Access, vol. 5, pp. 3157–3171, 2017.

[21] M. M. Sorkhabi and et al., “Measurement of transcranial magnetic stimulation resolution in 3-D spaces,” Measurement, vol. 116, pp. 326–340, 2018.

[22] R. Muhammad H., Power Electronic Circuits, Devices, and Applications, University of West Florida. Pearson Prentice Hall, 2004.

[23] J. I. Leon and et al., “The Essential Role and the Continuous Evolution of Modulation Techniques for Voltage-Source Inverters in the Past, Present, and Future Power Electronics,” IEEE Transactions on Industrial Electronics, vol. 63, no. 5, pp. 2688–2701, January 2016.

[24] L. G. Franquelo, J. Rodriguez, J. I. Leon and S. Kouro, “The age of multilevel converters arrives,” IEEE Industrial Electronics Magazine, vol. 2, no. 2, pp. 28–39, June 2008.

[25] A. Marquez and et al., “Variable-Angle Phase-Shifted PWM for Multilevel Three-Cell Cascaded H-Bridge Converters,” IEEE Transactions on Industrial Electronics, vol. 64, no. 5, pp. 3619–3628, 2017.

[26] E. R. C. da Silva and et al., “Pulsewidth Modulation Strategies,” IEEE Industrial Electronics Magazine, vol. 5, no. 2, pp. 37–45, June 2011.

[27] M. Srndovic and et al., “Simultaneous Selective Harmonic Elimination and THD Minimization for a Single-Phase Multilevel Inverter With Staircase Modulation,” IEEE Transactions on Industry Applications, vol. 54, no. 2, pp. 1532–1541, 2018.

[28] E. Barbie and et al., “Closed-Form Analytic Expression of Total Harmonic Distortion in Single-Phase Multilevel Inverters With Staircase Modulation,” IEEE Transactions on Industrial Electronics, vol. 67, no. 6, pp. 5213–5216, 2020.

[29] Y. Zheng and et al., “Research on DC Protection Strategy in Multi-Terminal Hybrid HVDC System,” Engineering, 2021.

[30] M. Memarian Sorkhabi, K. Wendt, D. Rogers and T. Denison, “Paralleling insulated-gate bipolar transistors in the H-bridge structure to reduce current stress,” SN applied sciences,vol. 3, no. 4, pp. 1–8, 2021.

[31] L. Wang and et al., PID and predictive control of electrical drives and power converters using MATLAB/Simulink, John Wiley & Sons, 2015.

[32] B. Wu and M. Narimani, “Two-Level Voltage Source Inverter,” in High-Power Converters and AC Drives, Wiley-IEEE Press, 2017, pp. 93–117.

[33] B. Sahoo and et al., “Repetitive control and cascaded multilevel inverter with integrated hybrid active filter capability for wind energy conversion system,” Engineering Science and Technology, an International Journal, vol. 22, no. 3, pp. 811–826, 2019.

[34] A. Barker, C. Garnham and I. Freeston, “Magnetic nerve stimulation: the effect of waveform on efficiency, determination of neural membrane time constants and the measurement of stimulator output.,” Electroencephalogr Clin Neurophysiol Suppl, vol. 43, pp. 227–37, 1991.

[35] S. Nurmi and et al., “Trade-off between stimulation focality and the number of coils in multi-locus transcranial magnetic stimulation,” Journal of Neural Engineering, 2021.

[36] H. Sinisalo, J. Nieminen and R. Ilmoniemi, “Waveform simulation and pulse-width-modulation approximations with multi-locus tms,” Brain Stimulation, vol. 14, no. 6, p. 1634, 2021.

