## Supplementary material for "A Neurostimulator System for Real, Sham, and Multi-Target Transcranial Magnetic Stimulation": Constricted coil for real, sham, and multi-target stimulation

### Appendix A

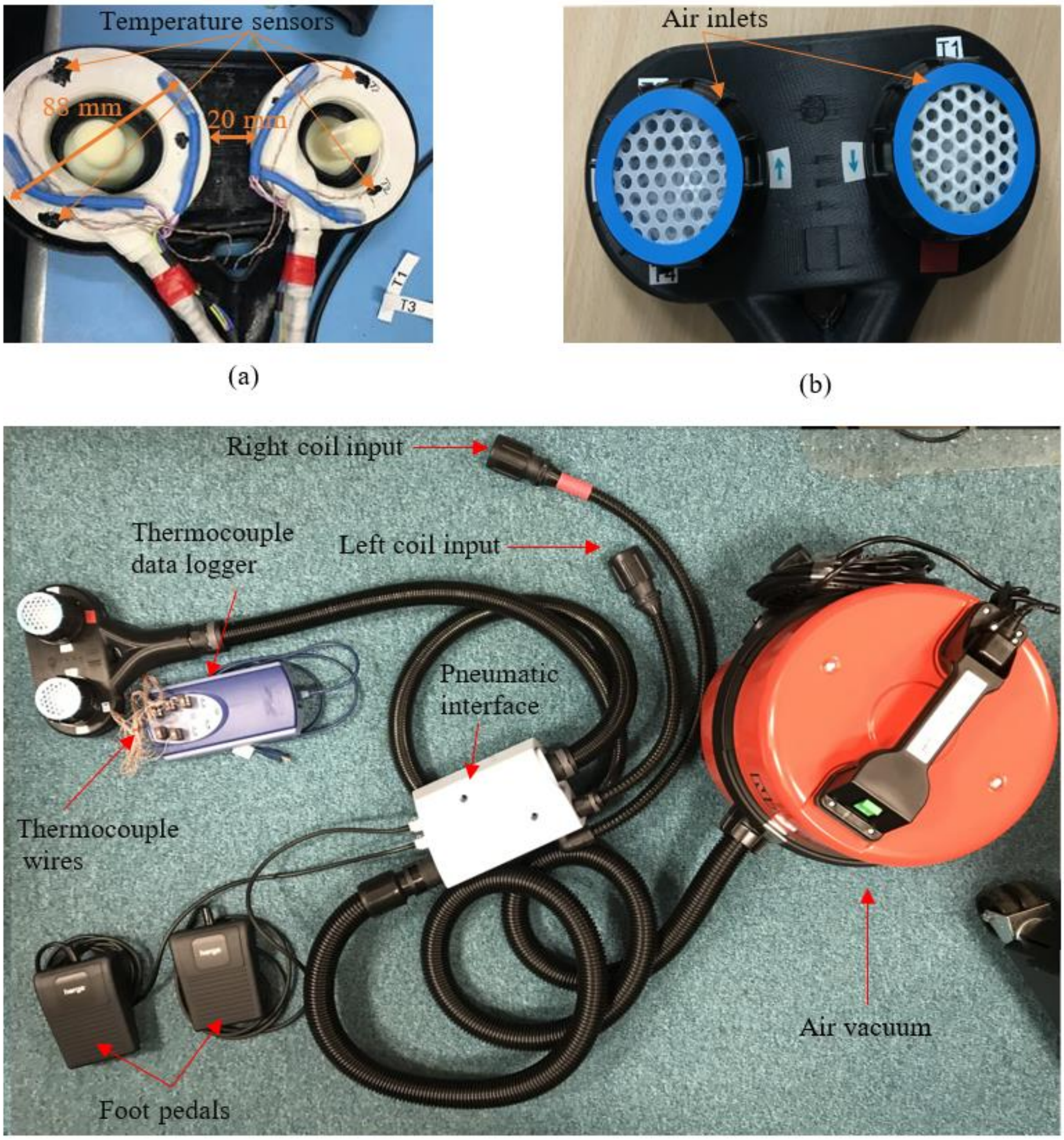


Figure A.1 Constricted coil for real, sham, and multi-target stimulation, (a) The coil consists of two circular coils with a physical distance of 20 mm between the coils, and four thermocouple sensors. Each coil consists of a tightly wound of 14 turns of wire with a cross-section of 10 mm x 1 mm. The inner and outer diameters of each coil are 52 and 88 mm, respectively. (b) The winding direction of each coil and two air inlets above the stimulation coil, for cooling. (c) Constructed coil detail, including high voltage coil inputs for the right and left coils, pneumatic foot pedals, thermocouple wires (type T thermocouple, RS Co., UK), thermocouple data reading equipment (8 channel thermocouple data logger, Pico Technology, UK), cooling system (air vacuum NQS 250-21, including audio noise reduction parts, Numatic Co., UK), and the pneumatic interface including pneumatic switches for the foot pedals and the air vacuum separator between coil inputs cables and the coil.
